# White matter microstructure characterisation in temporal lobe epilepsy

**DOI:** 10.1101/2021.05.08.442908

**Authors:** Nicolò Rolandi, Fulvia Palesi, Francesco Padelli, Isabella Giachetti, Domenico Aquino, Paul Summers, Elena Tartara, Giancarlo Germani, Valeria Mariani, Egidio D’Angelo, Laura Tassi, Claudia AM Gandini Wheeler-Kingshott, Paolo Vitali

**Affiliations:** Department of Brain and Behavioural Sciences, University of Pavia, Pavia, Italy; Neuroradiology, Fondazione I.R.C.C.S. Istituto Neurologico Carlo Besta, Milan, Italy; Neuroradiology, IRCCS Mondino Foundation, Pavia, Italy; Epilepsy Centre, IRCCS Mondino Foundation, Pavia, Italy; “C. Munari” Epilepsy Surgery Centre, ASST Niguarda, Milan, Italy; Brain Connectivity Center Research Unit, IRCCS Mondino Foundation, Pavia, Italy; NMR Research Unit, Queen Square MS Centre, Department of Neuroinflammation, UCL Queen Square Institute of Neurology, Faculty of Brain Sciences, University College London, London, United Kingdom; Radiology, IRCCS Policlinico San Donato, San Donato Milanese, Italy

## Abstract

Temporal lobe epilepsy (TLE) is the most common form of focal epilepsy. Parameters of microstructural abnormalities derived from diffusion tensor imaging(DTI) have been reported to be helpful in differentiating between Left and Right TLE (L-TLE and R-TLE) but few of them compared L-TLE and R-TLE with a voxelwise approach. In this study, a whole brain tract based spatial statistical analysis was performed on DTI, diffusion kurtosis and NODDI derived parameters of 88 subjects to identify specific white matter patterns of alteration in patient affected by L-TLE and R-TLE with respect to healthy controls. Our findings demonstrated the presence of specific patterns of white matter alterations, with L-TLE more widely affected both in cerebral and cerebellar regions. This result supports the need to consider patients separately, according to the side of their pathology.

## Introduction

Focal epilepsy is a chronic brain disease characterized by the occurrence of spontaneous and recurrent seizures [1]. These manifestations are characterized by clinical signs and symptoms connected with an abnormal neuronal activity manifested as hypersynchronous discharge of a population of brain neurons. The abnormal activity is transitory and difficult to predict, and symptoms depend on the brain area affected. Approximately 30-40% of patients with epilepsy shows drug-resistant epilepsy [2], meaning that seizures persist despite the administration of an adequate and well-tolerated antiseizures pharmacological treatment. Amongst all different types of epilepsy, temporal lobe epilepsy (TLE) is the most common form of focal epilepsy [3], [4]. The aetiology of TLE presents a wide variety of possible causes as shown by histopathological analysis, revealing combinations of focal acquired lesions, cortical malformations, hippocampal sclerosis[5]. Moreover, cognitive performance in subjects with epilepsy deteriorates frequently depending on the side of the epileptogenic zone; for example, a significant relationship between lateralized hippocampal pathology and memory dysfunction, i.e. verbal memory, is more evident in subjects with lesions in the left hippocampus [6]. A marked association between cognitive abilities and white matter (WM) abnormalities [7] has also been in TLE patients with the epileptogenic zone in the left hemisphere, whereas TLE patients with the epileptogenic zone in the right hemisphere mainly showed deficits in the recognition of emotional categories [8].

Magnetic resonance imaging (MRI) is routinely used to improve the diagnosis and to investigate the evolution of neurodegenerative pathologies and neurological diseases and can assume a crucial role for characterizing pathological conditions. Studies that have used MRI to detect alterations and to predict and investigate the surgical outcomes in epileptic patients, have revealed that TLE patients present structural abnormalities both in grey and white matter [9]. White matter abnormalities are generally considered secondary to the primary gray matter disease. Chronic degeneration has been demonstrated by one histological-7 MRI study on the resected tissue of the temporal pole in patients with hippocampal sclerosis [10] Another study using Diffusion Tensor Imaging (DTI) showed that white matter alterations are mainly ipsilateral to the seizure focus, but are more spread in TLE patients with the epileptogenic zone in the left hemisphere than in the right hemisphere[11], and controversial results have also been found in other DTI works [12],[13].

Diffusion-weighted MRI is widely used to study the microstructural architecture of the brain and its abnormalities. This characterization can be achieved because diffusion MRI is based on the investigation of the Brownian motion of the water molecules, which is hindered and restricted by the microstructural brain architecture, hence providing useful, though indirect, information about anisotropic properties of myelinated fiber bundles.

Diffusion tensor imaging (DTI) is the most basic application of diffusion MRI to provide quantitative invariant metrics, like the fractional anisotropy (FA), which are reproducible among subjects, sessions and scanners. More advanced diffusion models proposed over the years, have been demonstrated to be more sensitive and specific to microstructural alterations. Metrics defined using diffusion kurtosis imaging (DKI) [14], investigate diffusion proprieties beyond the gaussian behaviour captured by the DT, show greater sensitivity than DTI metrics in detecting and localizing changes in several diseases, [15][16], including TLE [17]. Efforts have also been dedicated to study the diffusion behaviour of water in brain tissue using computational models that depend on microstructural organisation assumptions. Among multi-compartment approaches, neurite orientation dispersion and density imaging (NODDI) is currently the most used, because it can be run on clinical scanners in relatively short acquisition times, is conceptually simple, and there are tools available for the analysis; indeed, NODDI is able to provide multiple parameters that can disentangle the different microstructural contributions to FA changes, such as the neurite density (ND) and the orientation dispersion (ODI) of dendrites and axons [18]. It has been proposed that this specific microstructural information is particularly useful in epilepsy because it could identify lesions such as cortical dysplasia that are often not recognized on other quantitative maps such as FA [19].

Global information about WM alterations can be provided by tract-based spatial statistics (TBSS), a fully automated data analysis technique that uses a voxel-wise approach to detect abnormalities in diffusion metrics, while minimizing the effects of misalignment introduced by registration in conventional voxel-based analysis methods [20]. The majority of TBSS studies in TLE patients, have investigated DTI metrics, while few studies have focused on DKI or NODDI-derived metrics. To the best of our knowledge, no study has investigated or reported differences between left and right lateralized TLE patients.

This work therefore, combines DTI, DKI, and NODDI-derived metrics on a large cohort of subjects, equally distributed in 3 different groups: healthy controls (HC), left TLE patients (L-TLE) and right TLE patients (R-TLE). TBSS was applied to DTI, DKI and NODDI-derived metrics in order to assess whether (i) specific patterns of WM alterations exist in left and right TLE groups, and whether (ii) DKI and NODDI-derived metrics provide information that is complementary to standard DTI-derived metrics.

## Methods

### Subjects

TLE patients were recruited as part of a project on 3T MRI in TLE subjects (the 3TLE project) from Epilepsy Surgery Centre ASST Ospedale Niguarda. The study was carried out in accordance with the Declaration of Helsinki with written informed consent from all subjects. The protocol was approved by the ethic committee of San Raffaele Hospital, Milan.

The inclusion criteria were diagnosis of temporal lobe epilepsy and potential candidate for surgery; resistance to antiepileptic drugs according to International League Against Epilepsy (ILAE) criteria (Kwan 2010) and age between 14 and 55 years. The exclusion criteria were large gliotic-malacic brain lesions and severe encephalic lesions; impossibility to perform MRI protocol due to physical or mental limitations, or contraindications to MRI.

TLE was diagnosed according to criteria defined by the ILAE [21]. All patients underwent a comprehensive neurological evaluation. Seizures were lateralized according to medical history, neurological examination, interictal electroencephalography (EEG), video-EEG, and in some cases, invasive recordings (Stereo-EEG), positron emission tomography (PET) and, after surgery, histological specimens. The correct identification of the epileptogenic temporal lobe is confirmed by the fact that temporal lobectomy in all cases leaded to significant seizure reduction, in most cases complete healing (class Ia, according to Engel classification of post-surgical outcome).

Age and gender matched healthy controls (HC) were also recruited from university students and hospital staff. The inclusion criteria were age between 19 and 55 years. The exclusion criteria were presence of current or previous neurological or psychiatric pathology, presence of two or more factors of cardiovascular risks.

### MRI acquisition

MRI data were acquired with a 3T Siemens Skyra scanner (Siemens Healthineers, Erlangen, Germany) using clinical and advanced sequences. Diffusion-weighted (DW) images were acquired with a two-shell twice refocused SE-EPI sequence using the following parameters: TR = 8400 ms, TE = 93 ms, 70 axial slices with no gap, 2.2 mm isotropic voxel, 45 volumes with non-collinear diffusion directions with b = 1000/2000 s/mm^2^ and 9 non-DW volumes (b = 0 s/mm^2^).

### DW analysis

Standard pre-processing steps comprising of correction for gibbs-ringing artifacts, noise floor, eddy-current induced geometrical distortions, and motion were performed on DW images combining commands from the MRtrix3 (https://www.mrtrix.org/) [22] and FSL (https://fsl.fmrib.ox.ac.uk/fsl/fslwiki/FSL) [23] toolboxes.

The pre-processed data were used to derive diffusion metrics under several models. DESIGNER (https://github.com/NYU-DiffusionMRI/DESIGNER) [24] was used to fit DTI and DKI and to calculate maps of FA, mean, axial and radial diffusivities (MD, AD and RD), and mean, axial and radial kurtosis (mK, aK and rK). NODDI provided maps of orientation dispersion index (ODI) and neurite density index (NDI).

### TBSS analysis

Whole brain voxel-wise statistical analysis was performed using TBSS (FMRIB, https://fsl.fmrib.ox.ac.uk/fsl/fslwiki/TBSS/UserGuide) [20]. FA maps from each subject included in the study (patients and HC) were non-linearly registered to each other in a process aimed at identifying the “most representative” subject of the group, which was then used as the target to align all FA images to the MNI152 (1×1×1mm^3^) template in standard space. All normalized FA maps were averaged and the mean FA image was thinned to create a mean FA skeleton of core WM voxels by imposing a threshold of FA=0.2. The normalized FA of each subject was projected onto the mean skeleton to create a subject-specific skeleton. The various skeletons were used as voxel-wise statistics to test for group differences. The other DTI, DKI and NODDI metrics were likewise projected onto the mean FA skeleton by applying the same non-linear registration used for projecting FA onto the MNI152 template.

The statistical voxel-wise analysis was performed with a permutation based inference approach of 5000 permutations on the skeletonized images for each metric using the randomize command (FSL). Age and gender were used as covariates. Comparisons were performed independently for all metrics (FA, MD, AD, RD, MK, AK, RK, ODI, NDI) using an ANOVA test. For each metric, results were corrected for multiple comparisons (family wise-error corrected) using the threshold-free cluster enhancement (TFCE) algorithm, and the significance level was set at p=0.05. Four pairs of groups were compared: 1) HC versus all-TLE patients; 2) HC versus L-TLE; 3) HC versus R-TLE and 4) L-TLE versus R-TLE.

### Temporal lobe Region Of Interest (ROI) analysis

A mask of the temporal lobes, i.e. a temporal lobe ROI, was extracted from an atlas of the brain lobes in MNI space. Mean values of DTI, DKI and NODDI derived-metrics were extracted for left and right temporal lobes as they are characterizing regions for TLE pathology.

### Statistical analysis

Statistical tests were performed using SPSS software version 21 (IBM, Armonk, New York). Demographic, clinical and diffusion-derived parameters were tested for normality using a Shapiro-Wilk test. Age was compared between groups (HC, L-TLE and R-TLE) using a two-tailed Kruskall-Wallis test while gender was compared using a chi-squared test. A Mann-Whitney test was performed on clinical features between the two groups of patients.

Since the three groups resulted to be gender-matched but not age-matched (HC vs R-TLE, p=0.009) a multivariate regression model with age and gender as covariates was used to compare all diffusion-derived metrics of the temporal lobe between groups, while Bonferroni correction was used to correct for multiple comparisons when comparing pairs of groups. Two-sided p < 0.05 was considered statistically significant.

A Spearman correlation analysis was performed between clinical features and diffusion-derived parameters of the temporal lobe in all patients, divided in left and right TLE.

## Results

### Patient Characteristics

Demographic and clinical features of all subjects are reported in table 1.

**Table 1:** Demographic table for Healthy controls and TLE patients.

|  | HC (n=36) | L-TLE (n=23) |  |  | R-TLE (n=29) |  |  | p-value |
| --- | --- | --- | --- | --- | --- | --- | --- | --- |
| Age (years) | 32.9 ± 8.4 | 33.1 ± 11.2 |  |  | 38.0 ± 9.9 |  |  | <0.033 |
| Gender (F/M) | 17 / 19 | 13 / 10 |  |  | 16 / 13 |  |  | n.s. |
| Age at onset (years) | - | 19.1 ± 11.8 |  |  | 23.8 ± 12.6 |  |  | - |
| Duration (years) | - | 14.0 ± 11.2 |  |  | 14.5 ± 11.6 |  |  | n.s. |
| Frequency (seizures/month) | - | 4.9 ± 6.2 |  |  | 6.6 ± 8.6 |  |  | n.s. |
| Histopathology / Engel Cls. | - | Tot | Ia | Other | Tot | Ia | Other |  |
| Cryptogenic | - | 4 | 2 | IIa, Ic | 15 | 9 | 6* |  |
| Hippocampal Sclerosis | - | 6 | 6 |  | 5 | 4 | Ic |  |
| Lesion | - | 9 | 7 | Ib, Id | 6 | 6 |  |  |
| Encephalitis | - | 1 | 1 |  | 1 | 0 | IVa |  |
| Polymicrogyria | - | 1 | 0 | IIa | 0 |  |  |  |
| Focal cortical dysplasia | - | 2 | 1 | Ic | 2 | 2 |  |  |

All values are reported as mean ± standard deviation. “n.s.” = not significant. In “Histopathology / Engel cls.” section:
Lesion = patients with tumor or cavernoma; Tot = total number of patients. Engel classification was obtained between
25-40 months after surgery. \* IIa, IIIa, IVa Ib e 2 with Id

Fifty-four TLE patients (35.8±10.7 years, 29 females) and 36 healthy controls (HC) (32.9±8.4 years, 17 females) were analysed. According to the criteria defined in the methods, twenty-three patients had the epileptogenic zone in the left hemisphere (33.1±11.2 years, 13 females) while 29 had the epileptogenic zone in the right hemisphere (38.0±9.9 years, 16 females).

For each patient group, Table 1 reports the histopathological classification according to the outcome of histological specimens and the Engel classification.

### Pattern of alteration in TLE patients

A first comparison was performed between HCs and all TLE patients in order to give a global description of WM alterations induced by TLE. Alterations were found through the brain in all diffusion metrics, except for AD and AK, as reported in Table 2.

**Table 2:**
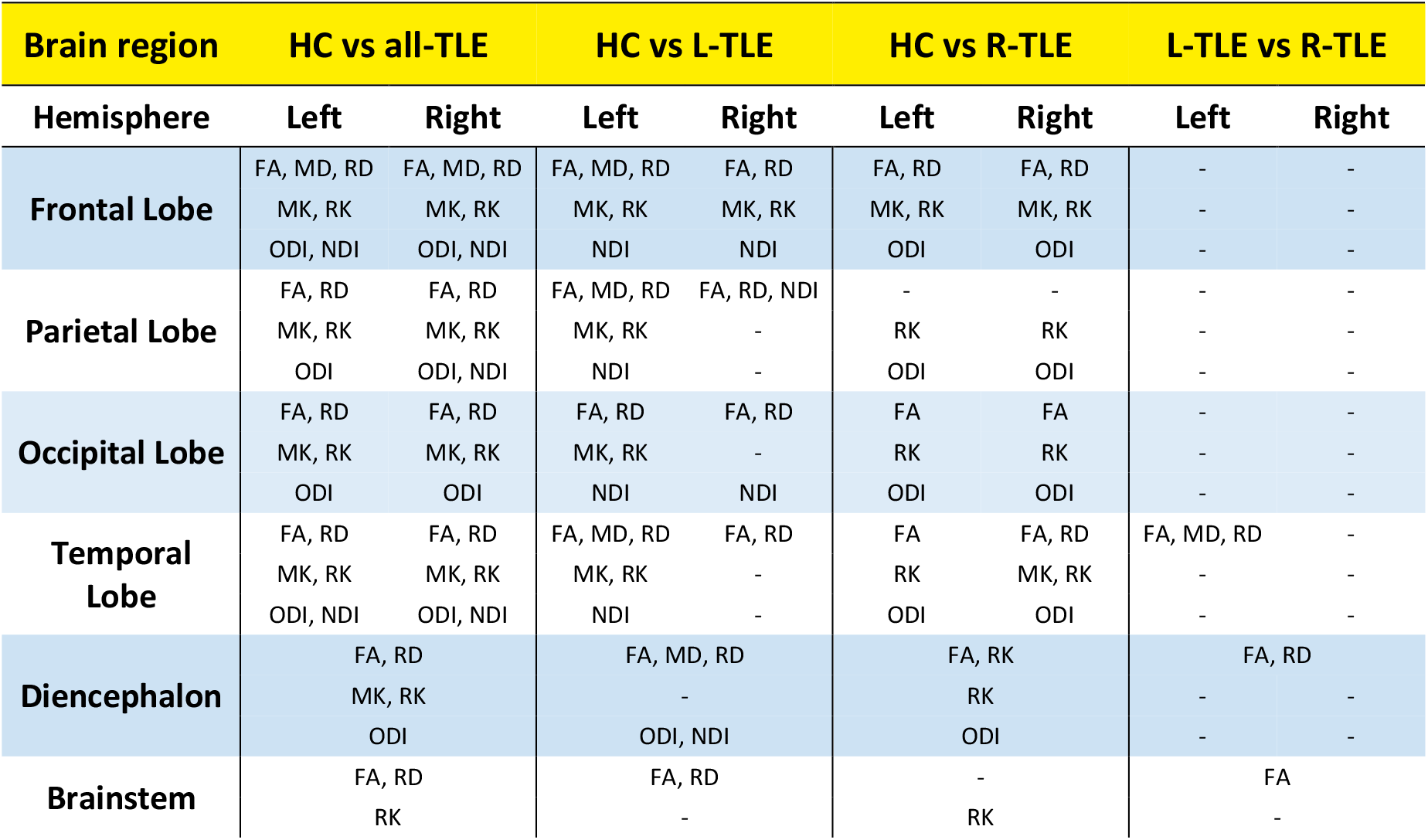
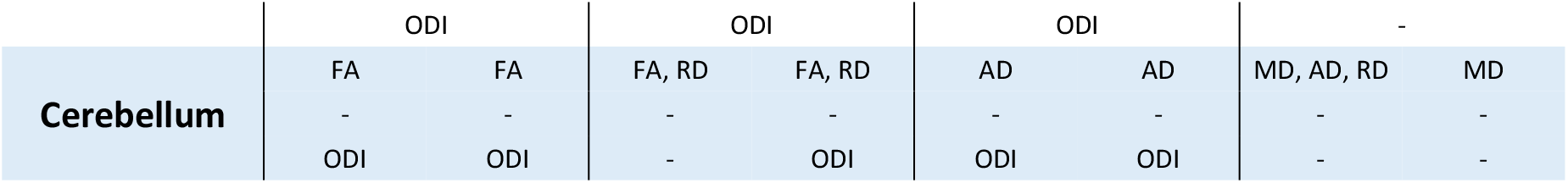
Metrics revealing significant alterations for each anatomical region in all TLE patients (all-TLE, second column), left TLE (L-TLE, third column), right TLE (R-TLE, fourth column) with respect to healthy control (HC) and L-TLE compared to R-TLE (fifth column). FA: Fractional Anisotropy; MD: Mean Diffusivity; AD: Axial Diffusivity; RD: Radial Diffusivity; MK: Mean Kurtosis; AK: Axial Kurtosis; RK: Radial Kurtosis; ODI: Orientation Dispersion Index; NDI: Neurite Density Index.

### Pattern of alteration in Left TLE

TBSS results on DTI-derived metrics revealed significant widespread bilateral FA reduction and RD increase in cerebral and cerebellar WM in L-TLE compared to HCs, with the exclusion of the right temporal lobe. Increased MD was predominantly found in the left cerebral hemisphere and involved diencephalon, temporal, parietal and frontal lobes (Figure 1a).

**Figure 1:**
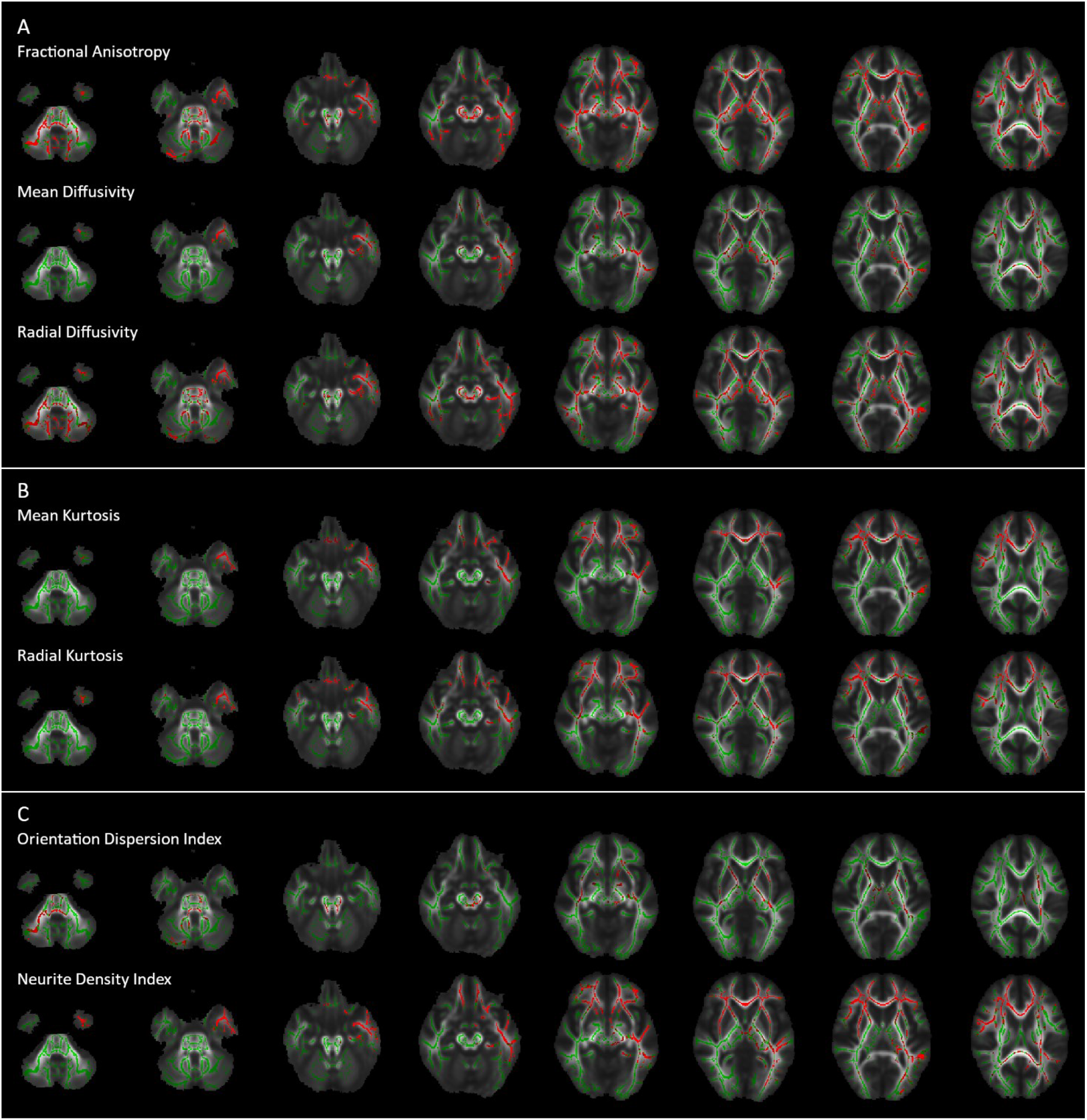
Alterations of maps in left TLE patients (L-TLE): (A) DTI, (B) DKI, (C) NODDI. L-TLE alterations are reported in red and the mean skeleton in green. Images are overlaid to the mean FA in MNI152 space, in axial radiological view.

DKI maps revealed decreased MK and RK in widespread brain regions, with the exclusion of the cerebellum, brainstem and a large portion of the right temporal lobe (Figure 1b).

NODDI maps showed widespread increased ODI in diencephalon, brainstem and right cerebellum. Changes in the temporal lobe were detected bilaterally, but were more pronounced in the left hemisphere, in the inferior longitudinal and uncinate fasciculi. Regions with decreased NDI involved many regions of the brain, predominantly in the left hemisphere, with the exception of brainstem and left hippocampus (Figure 1c).

### Pattern of alteration in Right TLE

R-TLE patients showed bilateral decreased FA in temporal, frontal and occipital lobes and diencephalon compared to HCs. Decreased AD affected the cerebellar peduncles and white matter bilaterally. Increased RD was mainly found in the right temporal and frontal lobes, plus the corpus callosum (Figure 2a).

**Figure 2:**
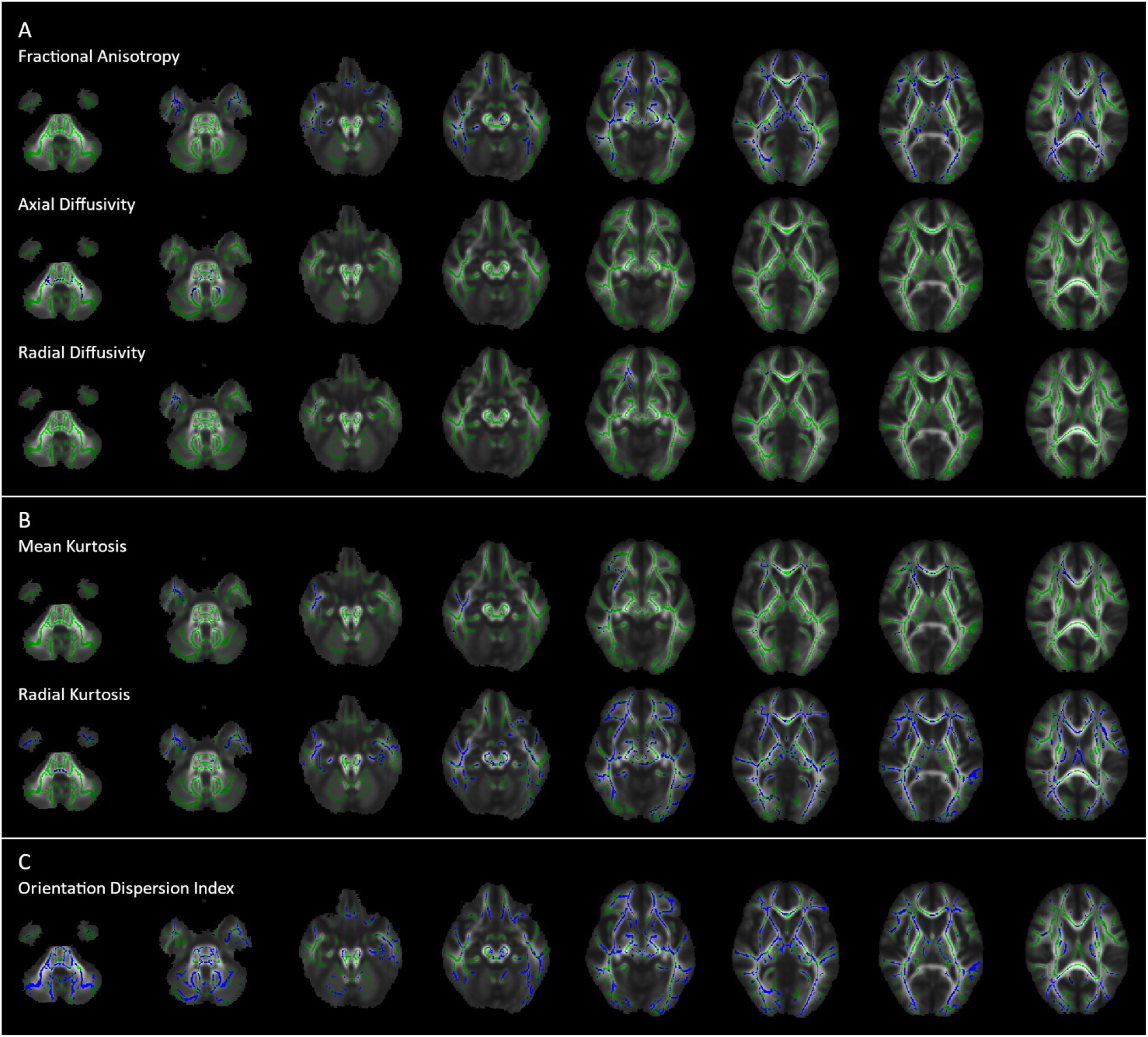
Alterations in right TLE patients (R-TLE): (A) DTI, (B) DKI, (C) NODDI. R-TLE alterations are reported in blue and the mean skeleton in green. Images are overlaid to the mean FA in MNI152 space, in axial radiological view.

**Figure 3:**
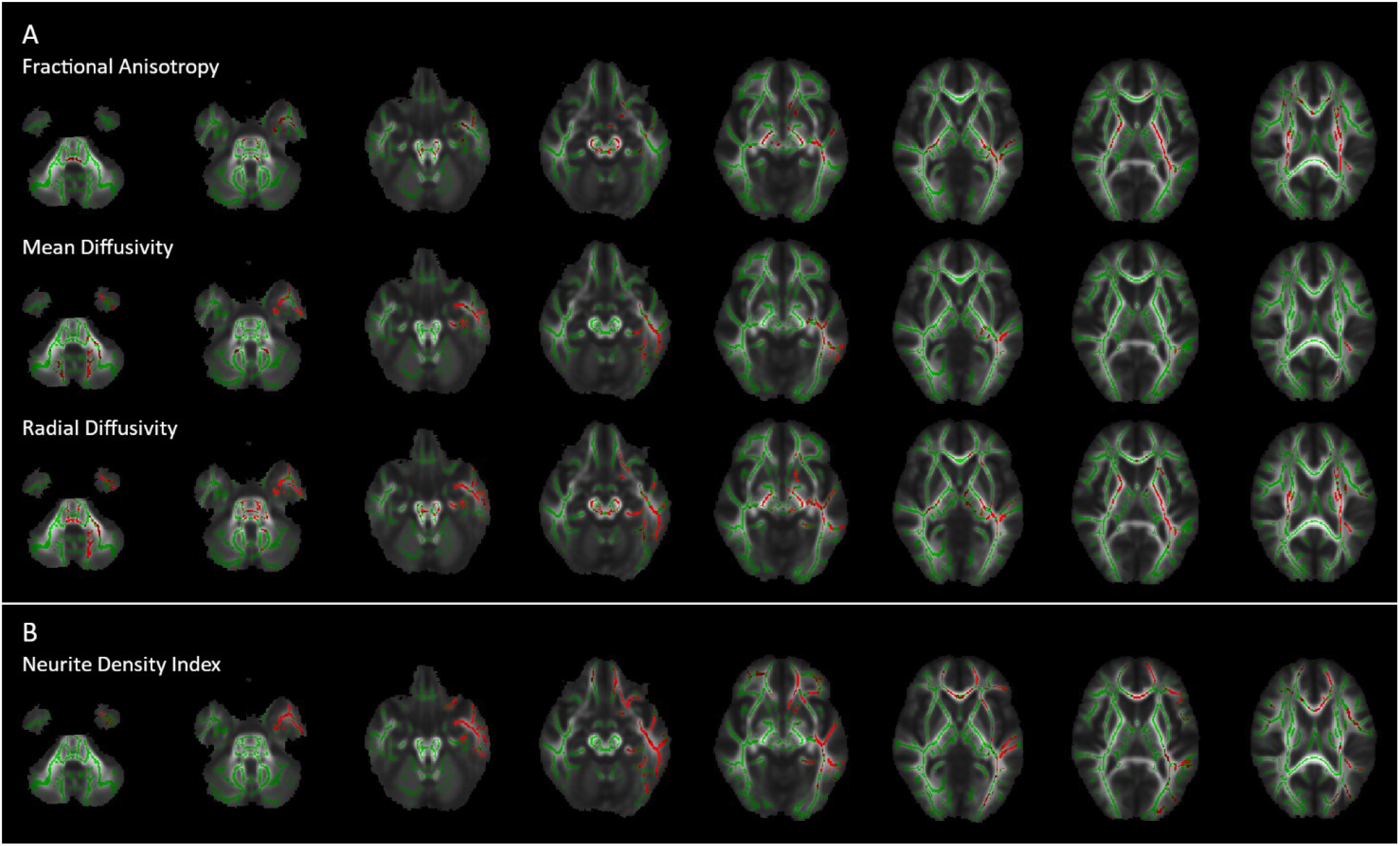
Alterations between patient groups: (A) DTI, (B) NODDI. L-TLE decreases compared to R-TLE are reported in red and the mean skeleton in green. Images are overlaid to the mean FA in MNI152 space, in axial radiological view.

Alterations in the right temporal lobe and bilaterally in the frontal lobe were revealed by decreased MK, while RK showed widespread decrease thought the brain (Figure 2b).

Increased ODI was widespread in the brain, but no NDI alterations were detected (Figure 2c).

### Differences between pathological groups

Compared to R-TLE patients, L-TLE patients presented decreased FA in the left temporal lobe, diencephalon and along the corticospinal tracts (CST), bilaterally. An MD increase was found in the left temporal lobe and rostrally along the superior longitudinal fasciculus (SLF), and bilaterally in the cerebellum. Increased RD was found in the left hemisphere, affecting both cerebrum and cerebellum, with particular involvement of the left CST and SLF.

R-TLE compared to L-TLE patients showed only decreased AD in the left cerebellum. Other metrics did not show significative differences.

### Temporal lobe ROI alterations

As shown in Table 3, in L-TLE compared to HC, the following significant differences between diffusion-derived metrics were found: decreased FA, MK and RK, increased MD and RD, decreased NDI in the left temporal lobe.

**Table 3:**
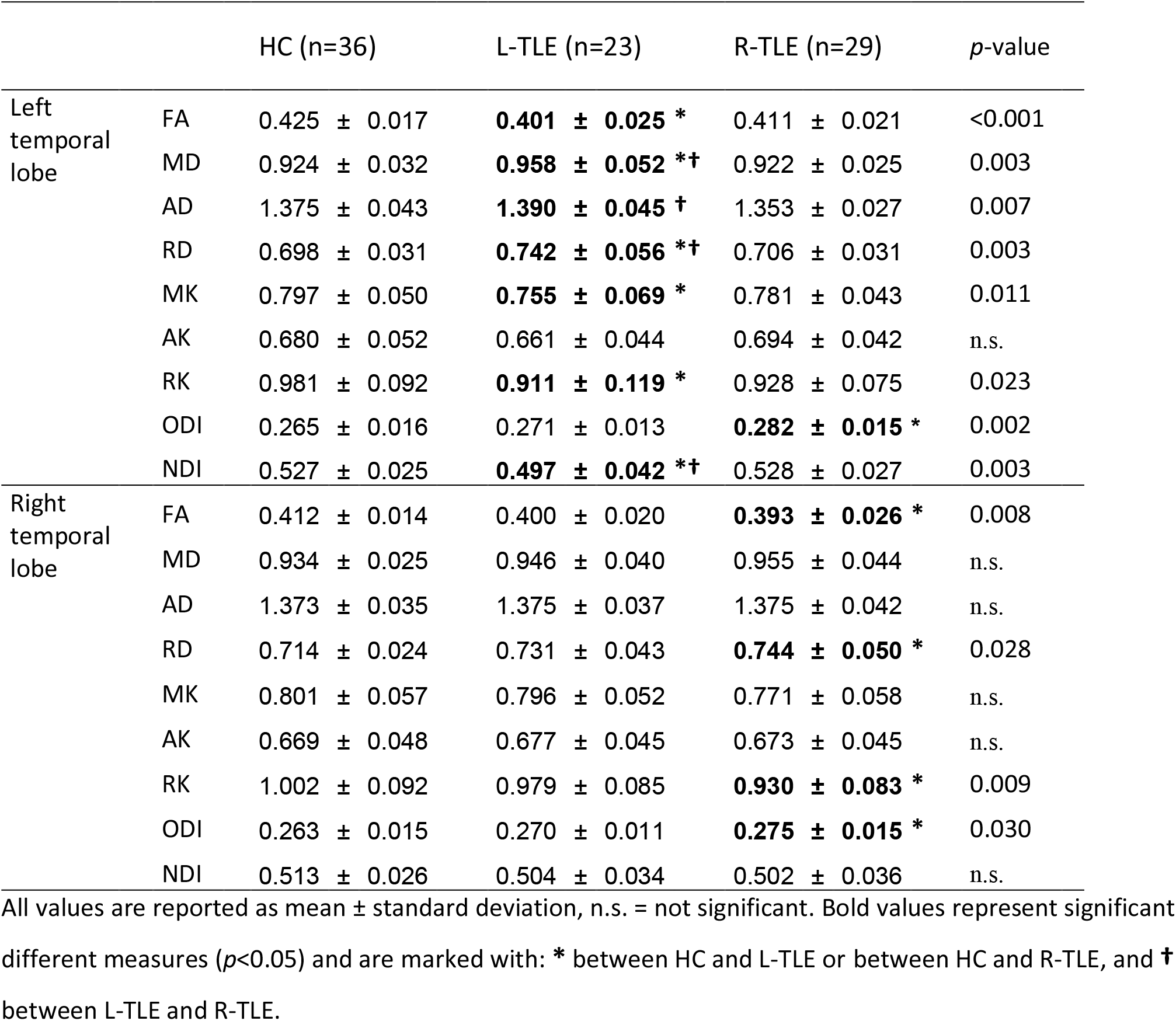
Temporal lobe alterations in TLE patients.

In R-TLE compared to HC, the following significant differences between diffusion-derived metrics were found: increased ODI in left temporal lobe, decreased FA and RK, increased RD and ODI In right temporal lobe.

In L-TLE compared to R-TLE, the following significant differences between diffusion-derived metrics were found: increased MD, AD and RD, decreased NDI in left temporal lobe

### Correlation between diffusion-derived metrics and clinical scores

In L-TLE, a significant negative correlation was shown between AK of the left temporal lobe and the frequency of seizures. In R-TLE, the frequency of seizures negatively correlated with MK and AK of the left temporal lobe, and negatively correlated also with FA, MK, AK, RK of the right temporal lobe.

No correlations were found between diffusion-derived metrics and illness duration.

## Discussion

The present work provides a novel characterization of TLE pathology. The main finding is that L-TLE and R-TLE present specific and not mirrored patterns of white matter microstructural alteration. Our findings revealed that the side of the epileptogenic zone is very important and determines consistent and specific microstructural changes. In particular, L-TLE involved more extended alterations than R-TLE in temporal and extratemporal areas, encompassing most of the brain. Indeed, here we have shown how different metrics help describing, and therefore may characterize structural alterations in a more marked way. Despite large inter-patient variability due to age, disease duration, and pathology underlying epilepsy, all patients present with drug-refractory epilepsy successfully treated by temporal lobectomy, which demonstrate that the epileptogenic area was confined within the resected temporal lobe. Therefore, all patients are clinically classified affected by TLE, with different underlying focal aetiology confirmed by histopathology. It was possible to report specific features of epilepsy and to investigate microstructural abnormalities interpreting them in relation to known biophysical changes. Most diffusion-derived indices revealed alterations in all TLE patients, independent on the side of epileptogenic zone, with respect to HCs. These widespread alterations, though, represent a “mean effect” of the pathology, hiding the presence of specific and focal alterations typical of left and right presentations, as demonstrated by our further results. If analysed overall, therefore, results could mislead interpretations. Being able to compare patients, grouped consistently according to clinical and physiological data, was necessary to identify underpinning important microstructural changes typical of a presentation of the disease; these could then become targets for intervention or could be used for assessing efficacy of interventions in clinical trials.

DTI and DKI demonstrated that L-TLE patients were characterized by a more extensive and generalized compromise of WM bundles with respect to HC. In particular, in L-TLE all DTI and DKI metrics revealed several bilateral regions of abnormalities and only MD was specifically increased in the left hemisphere. Interestingly, NODDI-derived metrics seems to be more sensitive and specific to microstructural alterations than DTI and DKI-derived metrics. Indeed, while DTI and DKI-derived metrics alterations, in particular FA ones, were widespread throughout the brain, NODDI-derived metrics demonstrated that such anisotropy alterations maybe due to different biological substrates as provided by NDI and ODI. These metrics, indeed, are complementary to each other and their spatial distributions overlap with all regions with reduced FA. In particular, a reduced NDI was detected only in cerebral regions in L-TLE patients, which is consistent with possible widespread WM alteration as shown in a previous work in epilepsy[25] and other neurological diseases[26], [27]. L-TLE patients also showed an increased ODI in brainstem and right cerebellum, indicating that here there might be a change in morphology of microstructure rather than cellular density [28]. Future post-mortem work should include histopathological analysis of cerebellar areas to confirm the interpretation of our findings. It would be interesting to run a longitudinal study to assess whether ODI changes precede NDI changes or whether these are stable and possibly linked to an underlying pathological malformation. Conversely, R-TLE patients showed more punctual and side specific alterations for all diffusion metrics compared to HC, suggesting a more localised pathology than in L-TLE. The most lateralized pattern was demonstrated by MK that was altered only in the right hemisphere. Again, it is interesting to look at results of NODDI-derived metrics, where ODI showed bilateral widespread alterations, while NDI was normal.

We can, therefore, hypothesis that cerebral and cerebellar microstructural alterations in L- and R-TLE might be characterised by different mechanisms, such as neurodegeneration or reduction of axonal density due to extracellular matrix changes. This difference could be also supported by the fact that the brain is functionally lateralized and functional reorganization due to pathology could be different. It is known that the left hemisphere is generally associated with language and analytic functions, the right hemisphere is mainly associated with visuospatial and global functions, while hippocampi are involved in the hippocampus the memory effect is lateralized to the left hemisphere, whereas navigation-related activity is prominent in the right hemisphere[29]. Furthermore, it has been demonstrated that in L-TLE language lateralization is decreased, and this correlate with language deficits before and after temporal lobectomy [30].This functional reorganization, predominant in L-TLE patients, and only to less extent in r-TLE patients, is likely to happen by reorganization of the WM structural connectivity respect to HC.

Longitudinal studies could help understand whether this ODI change precede an NDI alteration in the long term or whether it is a morphological abnormality typical of R-TLE epilepsy. Post-mortem studies are needed to underpin the biophysical substrate of this finding and clarify L-TLE and R-TLE WM differences. Although differences were found between L-TLE and R-TLE when comparing their advanced diffusion metrics from DKI and NODDI to those of HC, these metrics did not reach statistical significance when comparing L-TLE and R-TLE directly against each other. DTI metrics, though, revealed some differences, including the fact that the L-TLE group presented lower FA, and higher MD and RD in the left temporal lobe compared to the R-TLE one, consistent with the side of the more severe left pathology. Moreover, MD and RD showed specific alterations in the cerebellum, mainly in the left hemisphere. In our group we found lower AD in the left cerebellum among R-TLE respect to L-TLE patients. Given that the specificity of RD and AD to myelin and axon can be argued against[31], it is difficult to interpret these changes in terms of pathophysiology. Moreover, alterations are between two populations of patients rather than with HC and therefore they are indicative of specific pathological changes rather than the only changes.

From our results, DTI and DKI metrics can be very useful, showing different patterns of alteration of WM over the brain between L- and R-TLE respect to HC. Generalizing these results, we can assert that alterations are located mainly in the ipsilateral hemisphere of the epileptogenic zone and more widespread in the L-TLE group. Moreover, NODDI provides good discriminative pathological metrics between L- or R-TLE, especially when considering that NDI alterations have been found all over the brain in L-TLE while ODI all over the brain in R-TLE. From a brain network point of view, it would be interesting to localise such changes in terms of short and long-range connectivity, given that from a visual inspection of the results it seems that tracts like the corpus callosum and inferior fronto-occipital fasciculus are particularly involved.

Despite the fact that the cerebellum is often not mentioned in works on epilepsy when applying TBSS, studies with simultaneous EEG-fMRI was able to detect widespread functional alteration of cerebral networks with the involvement of the cerebellum in different type of focal epilepsy [32]. I It is worth noting that in our study the cerebellum was widely and differently involved in L- and R-TLE patients. In particular, FA reductions and RD increases were detected in the whole cerebellum of L-TLE patients, while AD and ODI increases were found in R-TLE. Moreover, cerebellar alterations were detected with anisotropic metrics (FA and ODI) or gaussian diffusion metrics (MD, RD, AD), while DKI metrics did not provide evidence of cerebellar impairment. The pathophysiological basis of these specific changes needs to be investigated in post-mortem studies and confirmed by independent cohorts of similar patients’ groups.

From ROI analysis in R-TLE patient group, emerged that only ODI was altered in the left temporal lobe. This is in agreement with TBSS findings, which suggest that NDI mainly reveals alterations in L-TLE while ODI is more specific for R-TLE alterations. Moreover, in left temporal lobe AD was significantly higher in L-TLE respect to R-TLE but no significant differences were found between the two pathological groups and HC. This result could suggest that while L-TLE show a compromission in the left temporal lobe, R-TLE present a functional reorganization in the healthy side.

It is important to note that, comparing TBSS and ROI results, several significant differences between groups were detected only with TBSS, which provides a global unsupervised information at voxel level, while ROI analysis, which gives only mean information in a specific brain region, might hide small focal and well-localized alterations.

Finally, interesting considerations could be made observing relations between diffusion-derived metrics and clinical features. First, only DKI metrics correlated with frequency of seizures. Moreover, while TBSS and ROI analysis revealed a greater compromised scenario in L-TLE, especially in the left temporal lobe, the correlation with clinical scores revealed a prevalent association between DKI metrics and frequency of seizures in the right temporal lobe of R-TLE. Despite these results are quite interesting, they must be carefully interpreted due to the high variability of the frequency of seizures among patients. Hence, further investigations are needed in order to improve knowledge regarding the physiological mechanisms underlying this correlation.

Despite our results being interesting for the specificity of the pathological changes due to TLE, this work represents only an initial evaluation of L-TLE and R-TLE specific patterns of alteration that should be improved by adding even more sensitive microstructural metrics, clinical and neuropsychological assessments. These improvements could help to investigate and predict surgical outcome and define possible cognitive rehabilitation therapy. In order to understand the underlying physiological and pathological mechanisms of TLE, multi-modal studies integrating imaging and data recording, e.g. from electroencephalography and functional imaging (PET and SPECT, MEG, EEG-fMRI), with cognitive and functional test are needed. These will also improve the monitoring and prediction of the pathological course in patients. Finally, the investigation of the relationship with clinical parameters, histological evidence and neuropsychological assessment of TLE patients is warranted in order to achieve a complete and comprehensive knowledge of the pathology.

## Conclusion

The present work reveals that L-TLE and R-TLE present different patterns of alteration, characterized by specific microstructural changes involving different brain regions, including the cerebellum. Compared to HCs, L-TLE patients are characterized by extensive and generalized damage of WM bundles, while R-TLE patients show more local WM alterations. The use of metrics from DKI and NODDI, other than FA from DTI, can help discriminating differences of WM alterations in sub-groups of TLE pathology. Despite these alterations can identify specific patterns for left and right TLE, the fact that quite all patients are seizure-free after surgery reveals that clinically relevant brain regions are localized near the epileptogenic zone. However, this suggests that the investigation of WM alterations spreading outside the temporal lobe might be important to understand specific mechanisms underlying TLE and its etiopathogenesis. Combining MRI microstructure assessment with neuropsychological and histopathological data will improve the characterization of TLE pathology and will provide not only possible outcome measures for clinical trials but also a mean to define more personalised interventions.

## Author Contributions

Conceptualized the study. designed and performed the analyses. provided support and guidance with data analysis and interpretation. coordinated the project and wrote the manuscript, with comments from all other authors.

## Acknowledgments

We thank the patients, their families, all healthy volunteers for making this research possible.

